# Composition and acquisition of the microbiome in solitary, ground-nesting alkali bees

**DOI:** 10.1101/2020.11.23.395194

**Authors:** Karen M. Kapheim, Makenna M. Johnson, Maggi Jolley

## Abstract

Increasing evidence suggests the microbiome plays an important role in bee ecology and health. However, the relationship between bees and their bacterial symbionts has only been explored in a handful of species. We characterized the microbiome across the life cycle of solitary, ground-nesting alkali bees (Nomia melanderi). We find that feeding status is a major determinant of microbiome composition. The microbiome of feeding larvae was similar to that of pollen provisions, but the microbiome of post-feeding larvae (pre-pupae) was similar to that of the brood cell walls and newly-emerged females. Feeding larvae and pollen provisions had the lowest beta diversity, suggesting the composition of larval diet is highly uniform. Comparisons between lab-reared, newly-emerged, and nesting adult females suggest that the hindgut bacterial community is largely shaped by the external environment. However, we also identified taxa that are likely acquired in the nest or which increase or decrease in relative abundance with age. Although Lactobacillus micheneri was highly prevalent in pollen provisions, it was only detected in one lab-reared female, suggesting it is primarily acquired from environmental sources. These results provide the foundation for future research on metagenomic function and development of probiotics for these native pollinators.

## INTRODUCTION

Communities of bacterial symbionts play an important role in animal biology, but the factors that shape the composition and acquisition of the microbiome are known for relatively few species. Rapid advancement in microbiome research has demonstrated that bacterial symbionts can influence host nutrition^1–3^, immunity^3–5^, and behavior^6–8^. Thus, understanding the health, physiology, or evolutionary ecology^9^ of any given animal species is incomplete without knowledge of their associated microbes. Two important aspects of this relationship are (1) the composition of the bacterial community throughout the host lifecycle and (2) the factors that determine how this community is acquired and maintained^10^. Despite a rapid advancement in microbiome research, knowledge of the relationship between animals and their bacterial associates is limited to a relatively small proportion of host species, especially among bees.

Understanding bee-microbiome relationships is particularly important, because bees are critical pollinators in both agricultural and natural communities. There is accumulating evidence that the microbiome influences several aspects of pollinator health^11,12^. For example, the microbiome affects nutritional intake by regulating appetitive behavior^13^, aiding in digestion^12,14–16^, or preventing spoilage of provisions^17^. An intact microbiome can also protect bees against toxins^18,19^, pesticides^19^, pathogens^20–23^, and parasites^24–26^, presumably in part by activating the host immune system^5,27,28^. Most of these findings stem from research with honey bees and bumble bees. It is unknown if similar protective effects of the microbiome are conferred to host solitary bees, partly because many wild bees lack a strongly characteristic core microbiome^29–36^. Thus, understanding pollinator health as it pertains to the microbiome requires knowledge of the factors that shape the composition and acquisition of the microbiome in a diverse set of bee species.

The factors that contribute to microbiome diversity are highly variable. Bee microbiome composition can be influenced by the evolutionary history^37,38^ and ecology^39–41^ of the host species, as well as intraspecific variation stemming from differences in caste^42–44^, development stage^34,45^, diet^46–48^, and infection status^49^. Research with additional bee species is likely to yield further insights into inter- and intra-specific variation in the microbiome. For example, most of the >20,000 described species of bees are solitary, nest underground, and diapause as larvae^50–52^. Yet, none of the bees for which the microbiome has been studied fit this ecological niche. Here we fill this gap with a study of the composition and acquisition of the solitary alkali bee *(Nomia melanderi)*, which is an important native pollinator in the western U.S.

Alkali bees are solitary, ground-nesting bees native to semi-arid regions of the western U.S. In some parts of their range, alkali bees are managed for alfalfa seed pollination, where they are encouraged to nest in moist soil beds sealed with salted surfaces^53^. This management practice results in some of the largest aggregations of bee nests ever recorded (up to 5.3 million) and gives alkali bees the unique distinction as the world's only managed ground-nesting bee^54^. Although they are highly effective alfalfa pollinators, alkali bees are floral generalists throughout their range^55^. Some of the threats they face include microbial spoilage of brood provisions^53,56^, viral infections^57^, larval predators^58,59^, cleptoparasites^53,59^, vertebrate predators^53^, collisions with automobiles^53^, and pesticide exposure^60^. Although some managed nesting aggregations have persisted for over 60 years, they are subject to extreme fluctuations in population size. Historical records of the current study population suggest there have been repeated population crashes followed by rapid and sustained growth^53,54,61^, and population genetic analyses suggest effective population size has declined in the recent past^62^. Alkali bees are facultatively multivoltine throughout their range, but univoltine in the current study population. Mating occurs in the spring or early summer, when males and females who have overwintered as pre-pupae complete diapause and emerge from their natal nests^53,63,64^. Females excavate a nest tunnel and begin provisioning brood cells within a few days of emergence. Each female provisions 9-16 brood cells within a 4-6 week adult lifespan^53^. We characterized the community of bacterial associates of alkali bees throughout their lifecycle (Fig. 1), and experimentally investigated how the adult female microbiome is acquired by identifying bacterial taxa that are differentially abundant in newly-emerged, lab-reared, and wild nesting bees. Our results provide an important reference point in understanding the relationship between bees and their microbial symbionts.

**Fig. 1.**
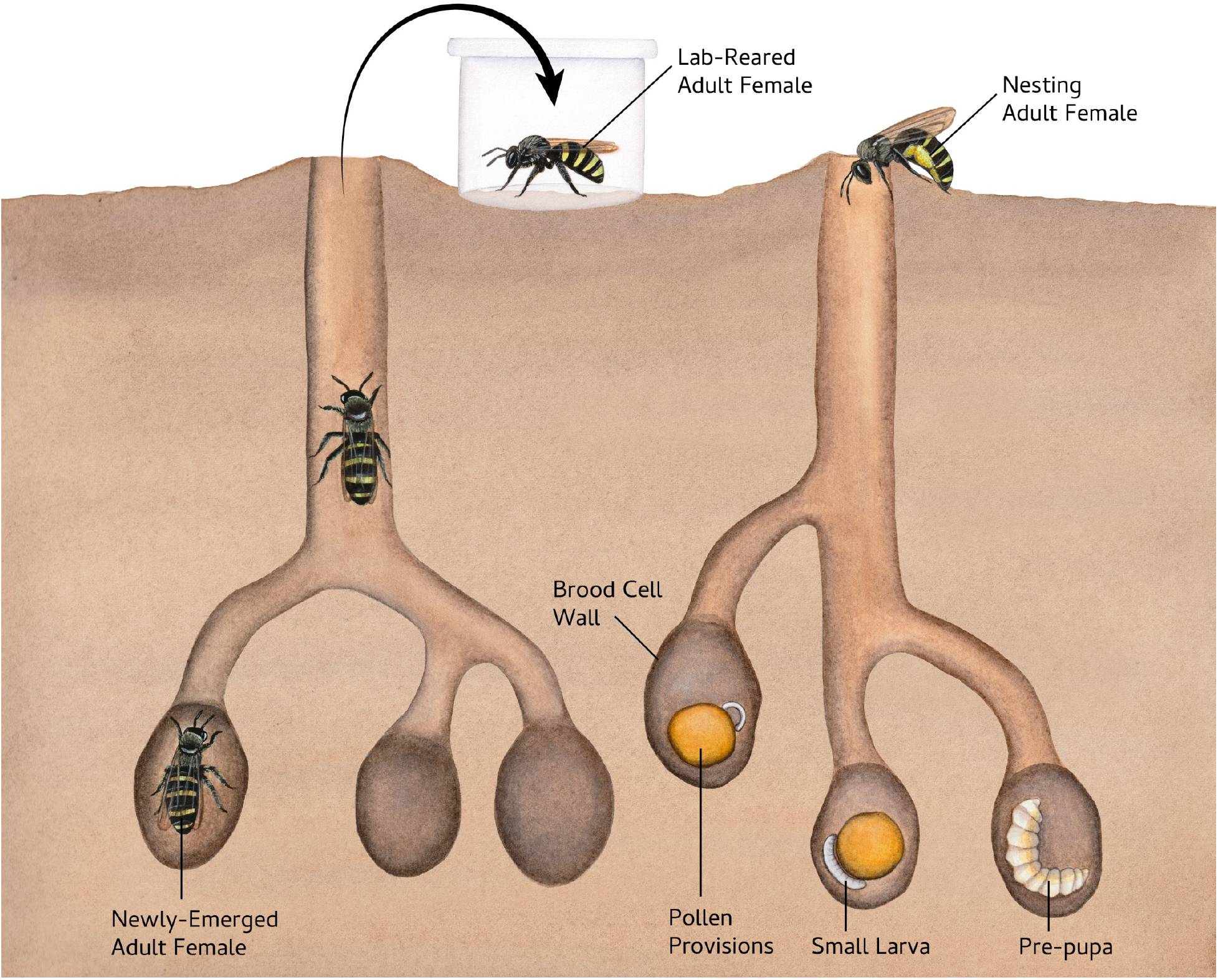
Experimental overview. We sampled adult females as they emerged from their nests after completing development in early summer. Some of these adult females were frozen immediately (newly-emerged) and others were reared in the lab for 10 d (lab-reared). We also sampled adult females that were free-flying and actively nesting (nesting females). We excavated nests to collect brood cell walls, pollen provisions, small larvae, and pre-pupae. Illustration by Julie Johnson (Life Science Studios).

## METHODS

### Bee Collections

Alkali bees *(Nomia melanderi)* were collected in June-July 2016 in Touchet, WA, USA. In Touchet, alfalfa seed growers maintain large beds of soil (called “bee beds”) that attract alkali bee nesting at very high densities^54^. We excavated nests in bee beds to collect uneaten pollen provisions from under eggs, small feeding larvae, pre-pupae (post-feeding larvae), and portions of the brood cell walls, which are lined with hydrophobic secretions in halictid bees. We used gloves and cleaned our tools with 10% bleach between each sample. Samples were transferred to clean 1.8 ml centrifuge tubes while in the field and frozen in liquid nitrogen within one hour of collection.

Adult females were the same as those used in a prior study^65^. We collected nesting females in nets returning to their nests with pollen on their legs, which indicates they were actively provisioning brood cells. Newly-emerged females were collected in emergence traps as they emerged from winter hibernation, following previously described methods^65^. Adult females were transferred to the lab in coolers and then either frozen in liquid nitrogen (newly-emerged and nesting females) or reared in the lab for 10 d under experimental conditions. Lab-reared females were randomly assigned to a diet treatment: sugar water only (sterile 35% sucrose solution), sugar water with pollen (2.5 g sterile, finely ground, honey bee pollen in 20 ml of sterile 35% sucrose solution), and sugar water with pollen plus four sprigs of fresh, un-tripped alfalfa flowers collected from fields adjacent to bee beds. Gamma-irradiated honey bee pollen was purchased from Better Bee. We pre-made individual 2.5 g packets of pollen with an additional round of sterilization via ethylene oxide (Anprolene AN74i), which were then vacuum sealed and frozen until use. Sterilization was confirmed by a lack of bacterial growth after plating and incubating a subsample of the sterilized pollen for > 72 hours. Sucrose solution was sterilized through a 0.2 micron filter following previous studies^66–68^. The pollen-sugar mixture was homogenized before each feeding and then pipetted into feeders. Fresh diet was prepared and feeders were cleaned with 10% bleach daily. Bees were maintained in plastic cages (72 mm x 90-113 mm) under full spectrum lighting (13 h light: 11 h dark) at 22-28 °C and 40-85% relative humidity. Cages were cleaned with 10% bleach prior to use. Upon collection, all samples were stored in liquid (or dry for shipping) nitrogen until return to Utah State University, where they were stored at −80 °C until dissection. Dissections followed previously reported methods^69^.

Importantly, the newly-emerged and lab-reared adult females were not “germ-free”. Each had some exposure to environmental sources of bacteria, but these differed from those of the freely nesting females, whom were actively foraging and were thus exposed to flowers and other elements of the landscape. Newly-emerged bees overwintered and completed development in the underground nests that they emerged from at the time of collection. They were thus exposed to bacteria present in the brood cell or nest tunnel and potentially to siblings who completed development at the same time. However, they did not have any exposure to the external environment (e.g., flowers), and they were prevented from interacting with other bees that had environmental exposure, because the traps prevented entry from the outside. Additionally, we aimed to eliminate bacterial inoculation from floral resources by pre-sterilizing the pollen and sucrose solution provided to lab-reared bees, but the lab conditions were not themselves sterile. Thus, the lab-reared females were exposed to bacteria present in the lab, but were deprived of the type of environmental exposure adult bees experience under normal, nesting conditions (e.g., flowers).

### DNA Extraction

We extracted DNA from each sample using MoBio PowerSoil kits, following manufacturers protocol, but with the addition of a 10 min incubation at 95 °C immediately following the addition of C1 solution. Working areas were cleaned with 10% bleach prior to extraction, and tools were flame sterilized between each sample. We extracted DNA from the hindguts of adult females following dissection. Larvae and pre-pupae were surface sterilized in a 1% bleach solution, followed by 3 rinses in sterile water. For larvae and pre-pupae, a 2 mm^3^ section was excised from the posterior end for DNA extraction. For pollen provisions, a 2 mm^3^ piece was excised from the center of the provision. We included a blank in each batch of extractions to control for contamination. These 12 blanks were included in the library preparation, sequencing, and sequence processing. DNA was eluded in 100 μl of C6 buffer, and yield was quantified with a Qubit HS DNA assay.

### Sequencing

The V4 region of the 16S rRNA gene was amplified on a Fluidigm Access Array for amplicon sequencing. We used the primers 515F (5'-GTGYCAGCMGCCGCGGTAA) and 806R (5'- GGACTACNVGGGTWTCTAAT). The resulting library was quantified by qPCR and sequenced on one MiSeq flowcell for 251 cycles from each end of the fragments using a MiSeq 500-cycle sequencing kit (v2). Fastq files were generated and demultiplexed with the bcl2fastq (v2.17.1.14) conversion software (Illumina). This generated a total of 20,024,886 reads from 84 experimental samples, with a mean ± standard error of 238,391.50 ± 27,122.47 reads per sample. Library preparation and sequencing were performed by the Keck Center for Comparative and Functional Genomics in the University of Illinois Biotechnology Center.

### Sequence Processing

After visually inspecting the distribution of quality scores, we processed the 16S rRNA sequences in the QIIME2 (v2019.4) environment. We used cutadapt to trim any remaining adapters. We then used DADA2 to join denoise and deplicate sequences, including the removal of chimeric sequences, singleton reads, quality filtering and joining of paired ends. We truncated forward reads at 213 nts and reverse reads at 191 nts, based on the location at which median quality score dropped below 30. We classified the resulting amplicon sequence variants (ASVs) with the SILVA 16S rRNA database (v132), using the 7 level taxonomy file and 99% identity. We extracted reference reads based on our 515F/806R primer pairs and length 100-400 nts. We then classified the ASVs with ‘classify-sklearn’. We aligned sequences with MAFFT and then generated a rooted phylogenetic tree FastTree using align-to-tree-mafft-fasttree. We then removed ASVs classified as mitochondria or chloroplast. We visualized rarefaction curves with ‘alpha-rarefaction’. Code is available at https://github.com/kapheimlab.

### Statistical Analysis

We performed statistical analysis of bee microbiomes in R v.3.6.2 ^70^, using the phyloseq v.1.28.0 tool ^71^. R code is available at https://github.com/kapheimlab. Two potential contaminants were identified and removed from the feature table with decontam v.1.4.0 ^72^, based on a criteria of being prevalent in more negative controls than real samples. We also identified two taxa that were detected in one negative control and one or more samples. This could be the result of tag-jumping, so we removed these taxa from samples for which the abundance was more than twice as high as it was in the negative control. This resulted in removal of one taxa from 9 samples. No samples met this criterion for the second taxa. We removed ASVs that were not assigned to a Phylum and which were not seen at least 25 times in at least 2 samples from the entire dataset. We also removed samples with fewer than 400 reads. Of the seven lab-reared females remaining in the dataset, two were fed only sugar water, four were fed sugar water with pollen, and one was given sugar water with pollen and fresh alfalfa sprigs. We removed the one lab-reared female given alfalfa. We then visually (Principal Coordinates Analysis [PCoA]) and statistically (adonis2 in Vegan^73^) investigated differences in the microbiome of lab-reared females given sugar or sugar and pollen. These two groups did not significantly differ (F = 1.33, d.f. = 1, p = 0.47; Fig. S1). We, therefore, collapsed these two sample types into a single ‘lab-reared’ category for all further analyses. Our final phyloseq object included 1,334 taxa and 62 samples. We rarefied to an even depth of 486 reads. Given the ongoing debate about the value of rarefaction^74^, we employed more than one normalization method where appropriate.

We visualized overall differences in microbial communities across sample types with Principal Coordinates Analysis (PCoA) applied to Bray-Curtis and weighted UniFrac distance matrices of log-transformed abundance data. We clustered samples with average linkage applied to a Bray-Curtis distance matrix of relative abundances. We tested for overall differences among sample types with adonis2 based on a Bray-Curtis distance matrix of relative abundances. We stratified 9,999 permutations across bee bed of origin. We followed this with pairwise comparisons using 9,999 permutation MANOVAs and a

Benjamini-Hochberg (BH) correction of p-values. We tested for differences in beta diversity with betadisper in vegan followed by pairwise comparisons with the Tukey Honest Significant Difference method (TukeyHSD)^75^.

We estimated the Shannon diversity index using a non-filtered dataset with the estimate_richness function in vegan. We tested for significant differences in square-root transformed Shannon indexes among sample types with a mixed effects model that included bee bed of origin as a random effect in the package lme4^76^. We used emmeans^77^ for pairwise comparisons with p-values adjusted by the Tukey method.

We used DESeq2^78^ to identify taxa that were differentially abundant across sample types. We included all sample types and bee bed of origin in the initial DESeq analysis, but then used pairwise contrasts to identify ASVs with differences in abundance that were significantly different at a BH-adjusted p-value < 0.05 between each type of adult female. Pearson’s correlations were measured between relative abundance of each ASV and metrics of reproductive physiology, including Dufour’s gland length, maximum terminal oocyte length, and maximum stage of oogenesis among adult females, using the associate wrapper in the microbiome package v.1.6.0^79^. We created a phylogenetic tree of ASVs classified as *Lactobacillus micheneri* with the function ‘plot_tree’.

## RESULTS

### Overall differences in microbiome

We identified significant differences in the overall microbial communities among sample types. PCoA revealed clustering among sample types (Fig. 2A). Specifically, Dimension 1 explained 22.9% of the variance in log-transformed microbiome composition and almost completely separated brood cell walls, pre-pupae, newly emerged females, and lab-reared females from pollen provisions, small larvae, and nesting females. This separation was also evident, though to a lesser degree, when the PCoA was based on a weighted unifrac distance matrix (Fig. S2) and in a dendrogram based on average-linkage of relative abundances (Fig. S3). Most sample types were dominant by bacteria from the phylum Proteobacteria, but the microbiome of pollen provisions and small larvae were comprised primarily of Firmicutes (Fig. 2B).

**Fig. 2.**
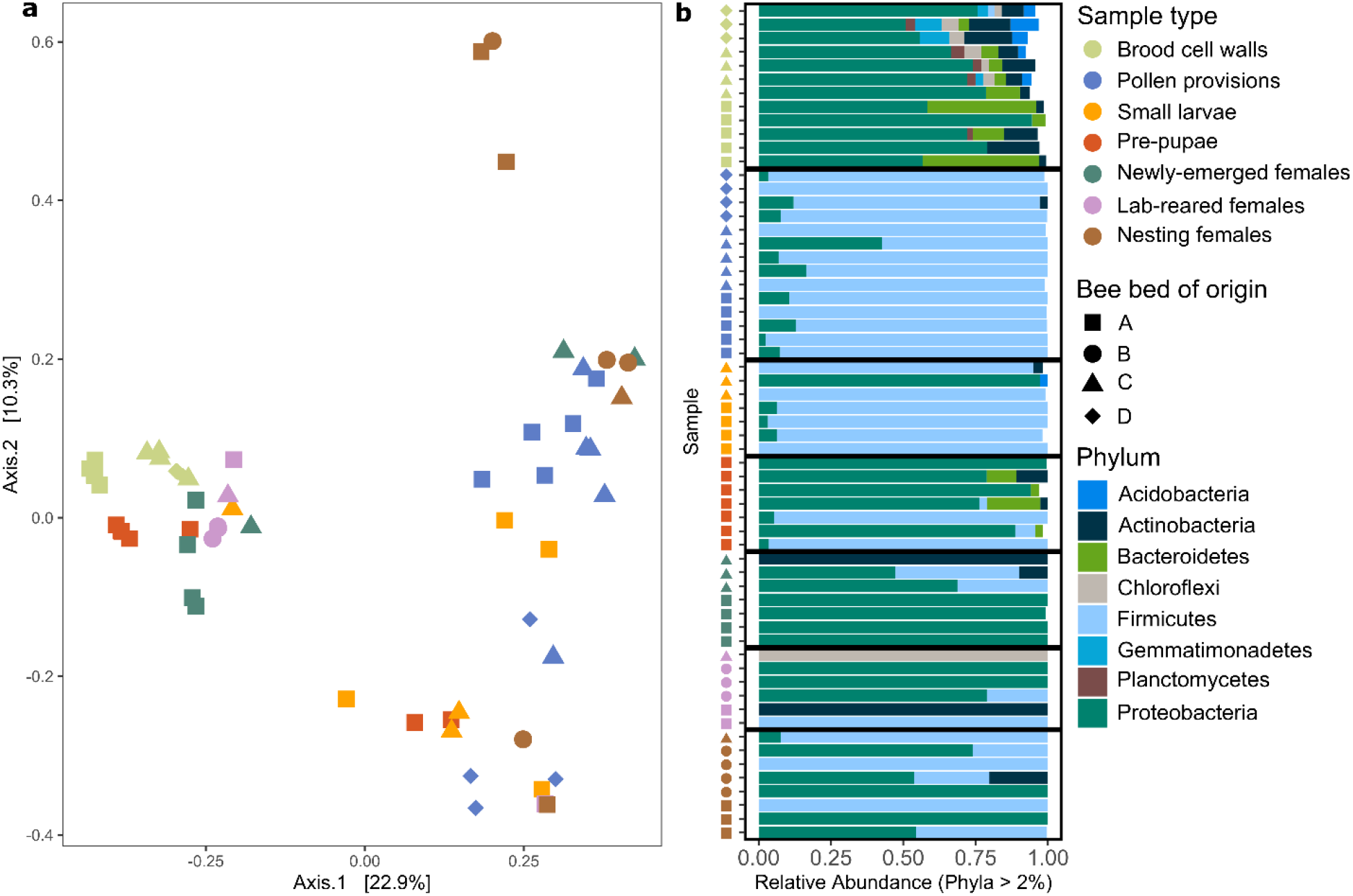
Microbiome composition across the alkali bee life cycle. (a) Principal Coordinates Analysis (PCoA) plot of Bray-Curtis dissimilarity from log-transformed abundances. Each point represents the bacterial community of an individual sample. (b) Relative abundance of Phyla found at greater than 2% abundance in each sample. Each row represents the bacterial community of an individual sample. Colors indicate sample type and shapes indicate bee bed from which sample was collected.

A permutation test revealed significant differences in the microbiome profiles among sample types (F = 5.356, d.f. = 6, p = 1e-04). Pairwise comparisons revealed significant differences (BH-adjusted p < 0.05) between all samples types except lab-reared vs. newly-emerged females (p = 0.076), pre-pupae vs. newly-emerged females (p = 0.052), and small larvae vs. pollen provisions (p = 0.135). Overall and pairwise results were consistent when this analysis was repeated on rarefied data (F = 5.537, d.f. = 6, p = 0.0001).

### Differences in diversity across sample types

There were significant differences in beta diversity, as measured by multivariate dispersion, among sample types (F = 6.444, d.f. = 6, p = 3.637e-05). Brood cell walls and adult females had the highest dispersion, while pollen provisions and small larvae had the lowest (Fig. 3A). Brood cell walls, newly-emerged females, and lab-reared females had significantly higher dispersion than pollen provisions and small larvae. No other groups had significant differences in dispersion. Overall and pairwise results were consistent when this analysis was repeated on rarefied data (F = 6.741, d.f. = 6, p = 2.281e-05).

**Fig. 3.**
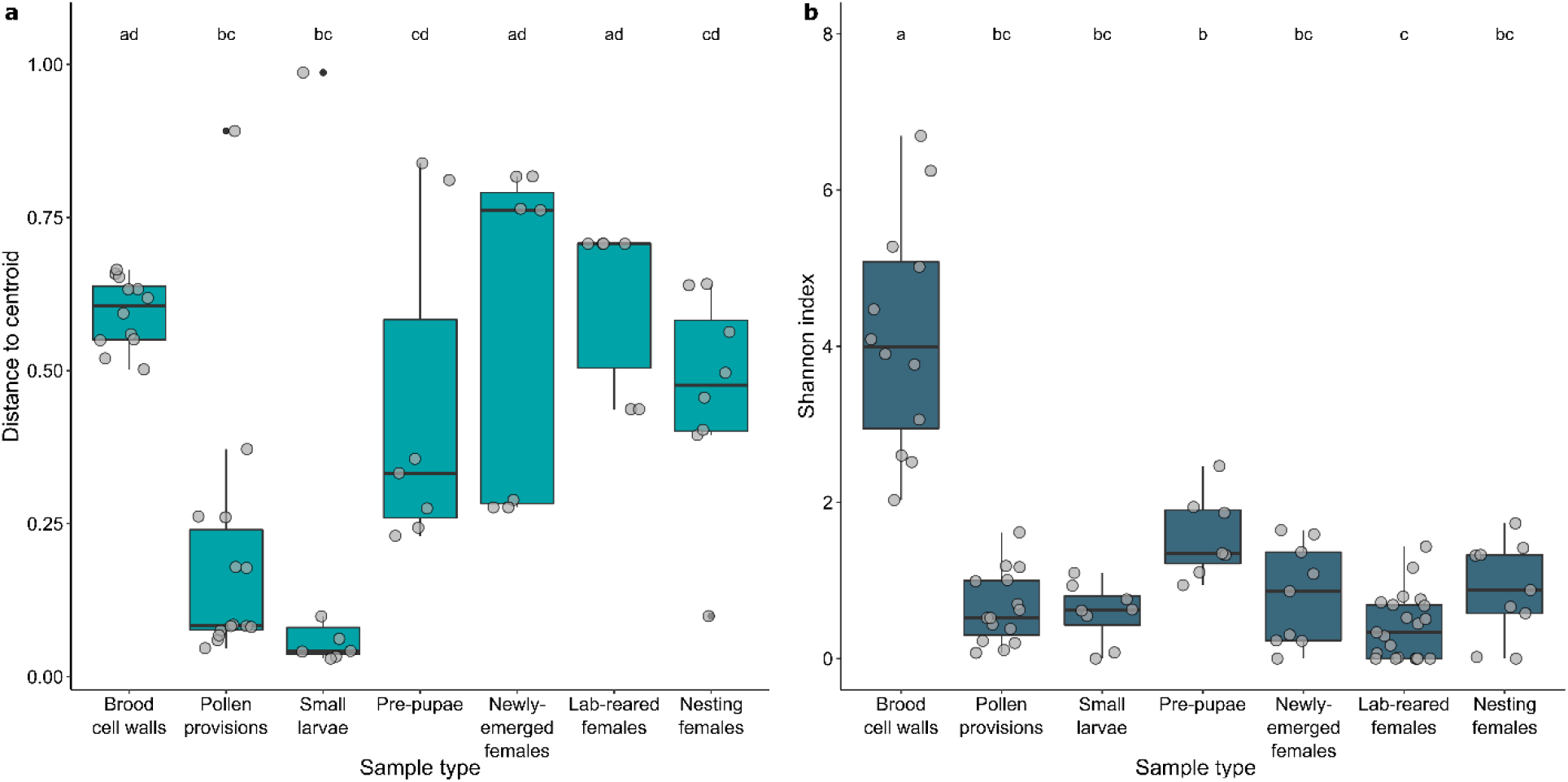
Bacterial diversity across sample types. (a) Multivariate dispersion displayed as distance from the centroid. (b) Alpha diversity calculated as Shannon index. Boxes represent the interquartile range, with a line indicating the median. Different letters along the top indicate significant (Tukey adjusted p < 0.05) differences between sample types. Whiskers extend to 1.5 times the interquartile values. Gray filled circles represent data from individual samples. There are more samples in (b), because unfiltered data was used to calculate Shannon index.

There were also significant differences in alpha diversity, as measured with the Shannon index, across sample types (F = 25.352, d.f., = 6, p < 3.415e-16; Fig. 3B). Brood cell walls had a significantly higher Shannon index than all other samples types (p < 0.003). Pre-pupae had a significantly higher Shannon index than lab-reared females (p = 0.006). These results were also consistent when the analysis was repeated on rarefied data (F = 28.856, d.f. = 6, p < 2.2e-16). However, in the latter case, pre-pupae also had a significantly higher Shannon index than small larvae (p = 0.03) and pollen provisions (p = 0.001).

### Differential abundance of key taxa and correlations

Overlapping sets of ASVs had significant differences in relative abundance between each type of adult female, and this allowed us to identify the potential source of each bacterial associate (Fig. 4; Table S1). ASVs that were significantly more abundant in the hindguts of nesting females than either newly-emerged or lab-reared females were likely primarily acquired from the external environment. Two ASVs met these criteria: one from the genus *Pseudomonas* and one classified as *Lactobacillus micheneri.* ASVs that were significantly less abundant in lab-reared females than in either newly-emerged or nesting females were likely acquired and maintained by contact with the nest environment. (Both newly-emerged and lab-reared females were exposed to the nest at emergence, but the lab-reared females could have lost these bacteria while kept in the lab for 10 d.) These included two ASVs from the family Enterobacteriaceae. We also identified ASVs for which relative abundance changed with age. Two ASVs had significantly higher relative abundance in newly-emerged females than both lab-reared or nesting females. These ASVs decreased in relative abundance with age and were classified as *Pseudomonas* and *Acinetobacter*. One ASV from the family Intrasporangiaceae increased in relative abundance with age (i.e., was significantly higher in relative abundance in lab-reared and nesting females than in newly-emerged females). We identified one ASV from the phylum Chloroflexi (soil bacteria Family JG30-KF-CM45^80^) that was likely associated with the lab environment, as it had significantly higher relative abundance in lab-reared females than in newly-emerged or nesting females. This pattern may have been driven by a single lab-reared female for which Chloroflexi dominated the gut microbiome (Fig. 2b). When this analysis was repeated with rarefied data, only *L. micheneri* was significantly more abundant in nesting females than in both lab-reared and newly-emerged females. No other taxa were significantly different between any groups of adult females. Correlation analysis failed to detect any ASVs that were significantly associated with Dufour’s gland length, maximum terminal oocyte length, or maximum stage of oogenesis (BH-adjusted p > 0.05).

**Fig. 4.**
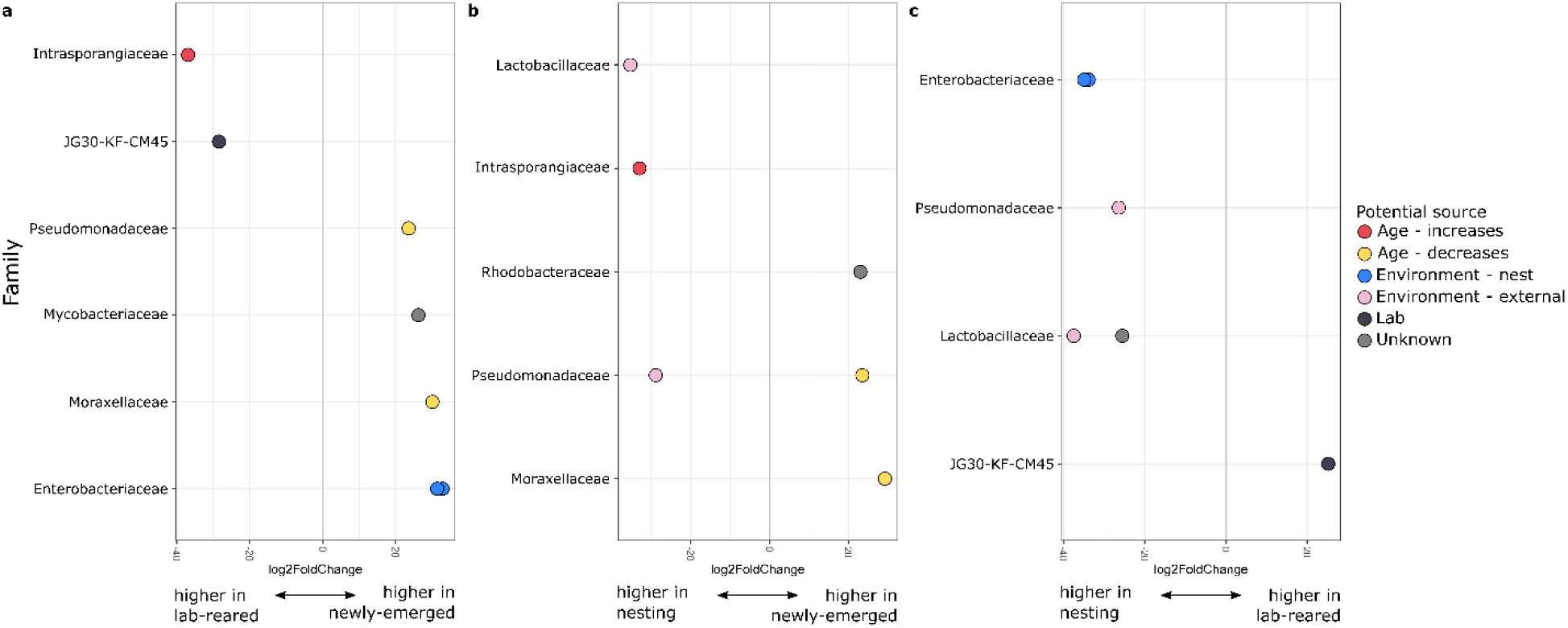
Differential abundance among adult females indicates potential sources of acquisition. Log2 fold change in hindgut relative abundance between (a) lab-reared and newly-emerged females, (b) nesting and newly-emerged females, and (c) nesting and lab-reared females. Filled circles represent a single ASV, with family membership indicated on the y-axis and color indicating potential source of acquisition.

### Lactobacillus micheneri

We further investigated the diversity and distribution of *Lactobacillus micheneri* among our sample types due to recent interest in how lactic acid bacteria are acquired in wild bees^32,33,40^. We detected 10 ASVs that were taxonomically classified as *L. micheneri. L. micheneri* has since been described as three distinct species - *L. micheneri, L. quenuiae*, and *L. timberlakei*^81^. It is therefore likely that many of these strains are actually different species. Indeed, the phylogenetic relationship of these ASVs reveals three main clades (Fig. 5). Many of these were specific to one or two sample types and at relatively low abundance (Table S2). One ASV was found in every type of sample, with the exception of brood cell walls. This was the only strain of *L. micheneri* that was detected in lab-reared females, and it was detected in all 14 of the pollen provision samples. *L. micheneri* was not detected in any of the brood cell wall samples, and was only detected in one of the six lab-reared female samples. *L. micheneri* diversity was highest among feeding larvae, as seven of the 10 ASVs were detected in small larvae. Six of the 10 ASVs were detected in the hindgut of nesting females. *L. micheneri* was relatively rare among newly-emerged females, with only three ASVs detected in one or two samples each.

**Fig. 5.**
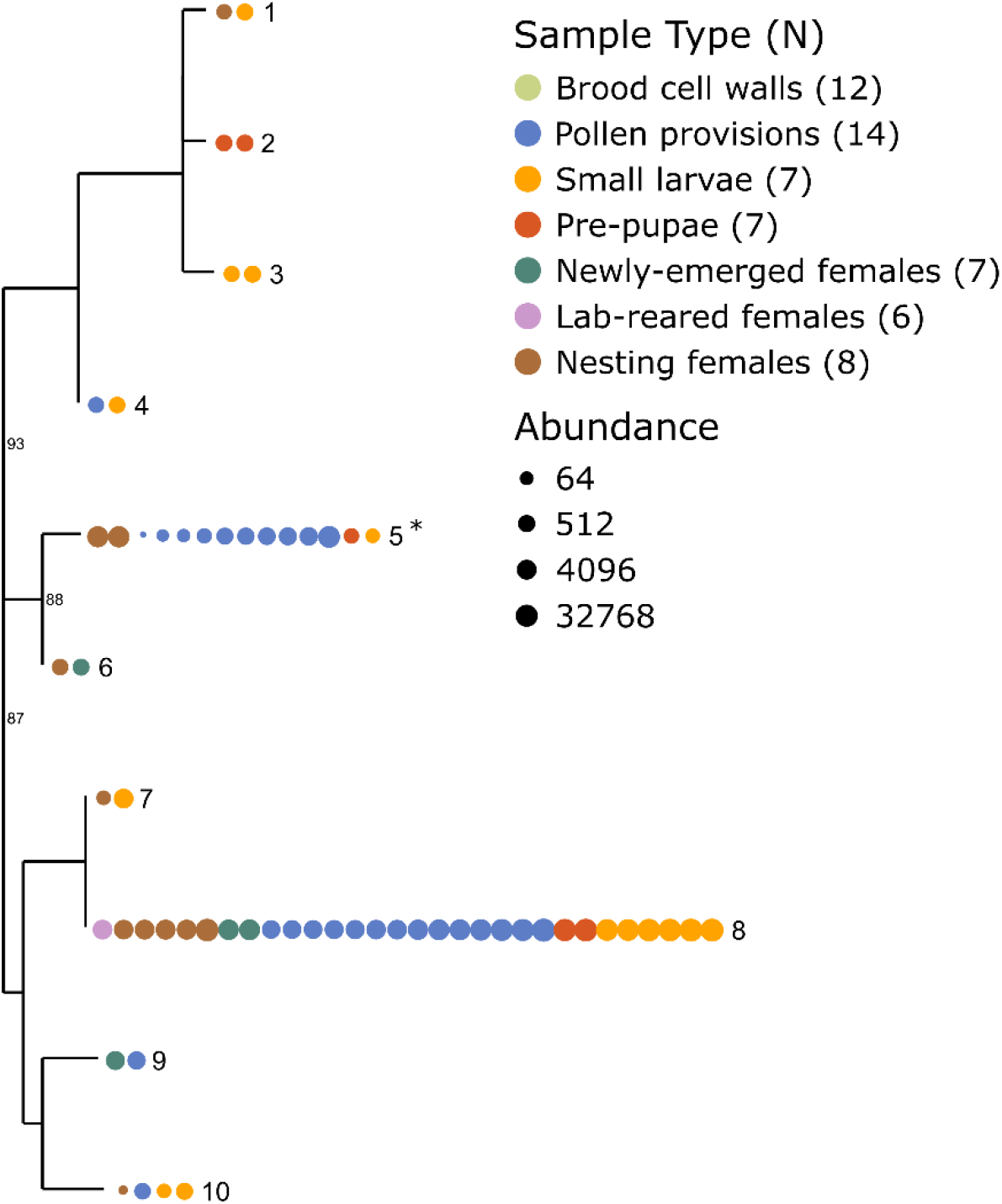
Diversity of *Lactobacillus micheneri* in alkali bees. Phylogenetic relationship of ASVs classified as *L. micheneri* with bootstrap support values near the nodes. Each circle represents an individual sample. Color indicates sample type and size reflects abundance on a log_2_ scale. Tip label identifies the ASV in Table S2. * signifies *Lactobacillaceae* identified as externally sourced in Fig. 4b,c. Numbers next to sample type indicate sample size (N).

## DISCUSSION

We characterized the composition and diversity of the alkali bee microbiome across its life cycle and experimentally investigated potential sources of key bacteria among adult females. Although this is the first description of the microbiome in a solitary ground-nesting bee, we find the most prevalent taxa are similar to those common in the microbiomes of other bees (e.g., Proteobacteria, Firmicutes)^82^. Our study shows that community composition of bacterial associates changes throughout the life cycle of alkali bees, as there were significant differences in the overall microbiome of feeding larvae, pre-pupae, newly-emerged, and nesting females. Comparisons of intra-group dispersion suggest most of these differences are not heavily influenced by differences in heterogeneity among our sample types. Moreover, we collected all samples at the same locations at the same time of year. Thus, overall differences in composition are not likely driven by seasonal or environmental fluctuations. Examination of microbial composition and diversity allowed us to make inferences about the factors that shape the microbial communities associated with alkali bees.

One of the clearest findings of our study is that the alkali bee microbiome is heavily influenced by feeding status. The community composition of bacteria found in the feeding larvae is highly similar to that of the pollen provisions collected from brood cells. This may reflect the fact that we sampled from the posterior (gut) end of the larvae, which was likely filled with recently consumed pollen. The hindguts of nesting females harbored a bacterial community that was also quite similar to that of feeding larvae and pollen provisions. (Adonis analysis revealed a statistically different community composition, but nesting females clustered with pollen provisions and small larvae on axis 1 of the PCoA plot.) Adult alkali bees regularly consume nectar and pollen^83^, so it is perhaps unsurprising that their gut microbiomes would be similar to those of brood provisions and feeding larvae. Yet, the significant difference in overall composition reveals there are likely to be unique resident bacteria living in the hindguts of adult female alkali bees. Nesting females have a relatively higher proportion of Proteobacteria from the family Enterobacteriaceae than small larvae and pollen provisions, which tend to be dominated by Firmicutes. Other members of the Enterobacteriaceae family found in honey bee guts *(Gilliamella apicola* and *Frischella perrara)* aide in digestion and immunity^12,84^, but it is unknown if the bacteria detected in alkali bees play similar roles.

Non-feeding larvae (pre-pupae) are more influenced by their environmental surroundings. Evidence for this is that the microbiome composition of pre-pupae and newly-emerged adult female hindguts were not significantly different. The primary source of contact for both pre-pupae and newly-emerged adults is the nest. Indeed, both of these sample types clustered with brood cell walls on axis 1 of the PCoA (Fig. 2A). This could reflect the fact that their guts are empty. Larvae typically expel meconium after they have finished consuming their pollen provisions and do not eat again until after they complete development and emerge from their natal nest. While it is known that honey bees acquire their microbiome from the hive environment^68^, the external environment is thought to play a larger role in determining the solitary bee microbiome^32,33,85^. Our results indicate that this is specific to development stage, particularly with regard to feeding.

Patterns of diversity allow inferences about the functional role of the microbiome across the alkali bee lifecycle. For example, pollen provisions and small larvae had the lowest beta diversity of any group. This indicates that the brood provisions of alkali bees are highly uniform, which could suggest the microbiome has a functional role in preventing spoilage, digestion, or other processes important to the early stages of bee development. This is consistent with the high prevalence of *Lactobacillus* (primarily *L. micheneri)* in the pollen provisions and small larvae. *Lactobacillus* are commonly found in pollen provisions and larvae of other wild bees^32,33,35^. In honey bees, a diverse flora of *Lactobacillus* play a role in activating the immune response^5^, inhibiting pathogens^21^, and preventing spoilage in stored pollen^86^. Genomic analyses suggest bacteria in the *L. micheneri* clade may be capable of inhibiting spoilage-causing pathogens and aiding in digestion and detoxification of pollen^87^. This suggests that the uniformity of a *Lactobacillus* based microbiome in alkali bee pollen provisions and small larvae is an adaptation that ensures optimal nutrition for developing alkali bees.

Our study also provides some insight as to how alkali bees acquire their bacterial associates. Newly-emerged and lab-reared females had statistically similar communities of bacteria in their hindguts. Yet the hindgut microbiome of nesting females was statistically different in overall composition from either newly-emerged females or lab-reared females. This suggests that the microbiome is substantially influenced by bacteria acquired from the environment, as has been suggested for other wild bees^40^. Additional analyses revealed that at least two ASVs are significantly more abundant in nesting females than in newly-emerged and lab-reared females. This suggests they are likely acquired from the external (potentially floral) environment. Bacteria in the *L. micheneri* clade are commonly transmitted between flowers and wild adult bees^32,33,40^. One ASV classified as *L. micheneri* (tip 5 in Fig. 5) was not detected at all in newly-emerged or lab-reared females. It also was not detected in brood cell walls and only at low levels in small larvae and pre-pupae. It was, however, detected in relatively high abundance in nesting females and pollen provisions. This suggests this bacterium is common on flowers, and that nesting females are frequently re-inoculated as they forage.

## CONCLUSION

Our study provides the first description of a solitary, ground-nesting bee, which also happens to be a native pollinator of economic import in the western U.S.A. Alkali bees occupy the ecological niche most common to bees across the globe. Understanding the patterns of microbiome diversity and acquisition in this species may provide insights about the relationship between bees and their bacterial associates that apply to other species. These insights include the following: (1) Composition of the microbiome changes over the course of development, and is largely influenced by food intake. (2) The bacterial make-up of pollen provisions (and thus feeding larvae) is highly uniform and largely comprised of *L. micheneri*, suggesting a functional role in early development. (3) The gut microbiome of nesting females is largely acquired after completion of development, and the external environment is likely to be an important source in this process.

## DATA AVAILABILITY

All raw sequence data have been deposited in the NCBI SRA (BioProject PRJNA675403). Code and data are available at https://github.com/kapheimlab.

## ACKNOWLEDGEMENTS

We are grateful to Craig Huntzinger and Diana Cox-Foster who assisted with pollen sterilization. Emily Thomas assisted with DNA extractions. We thank John Dodd and Forage Genetics International for providing lab space and logistical support in Touchet, WA, USA. We thank Mike Ingham, Mark Wagoner, and Mike Buckley for access to their bee beds and bees. F. Dowsett provided valuable assistance in the field. F. K. Hunter and M. A. Hagadorn provided valuable feedback on an earlier version of this manuscript. Funding was provided by the USDA-ARS Alfalfa Pollinator Research Initiative and Utah Agricultural Experiment Station (Project 1297).

## AUTHOR CONTRIBUTIONS

KMK designed the experiment, performed the experiment, analyzed the data, and wrote the paper. MMJ and MJ performed the experiment, edited the paper, and approved the final manuscript. MMJ performed dissections and DNA extractions.

## ADDITIONAL INFORMATION

The author(s) declare no competing interests.

