## Supplementary material for "Composition and acquisition of the microbiome in solitary, ground-nesting alkali bees": Fig. S1, Fig. S2, Fig. S3

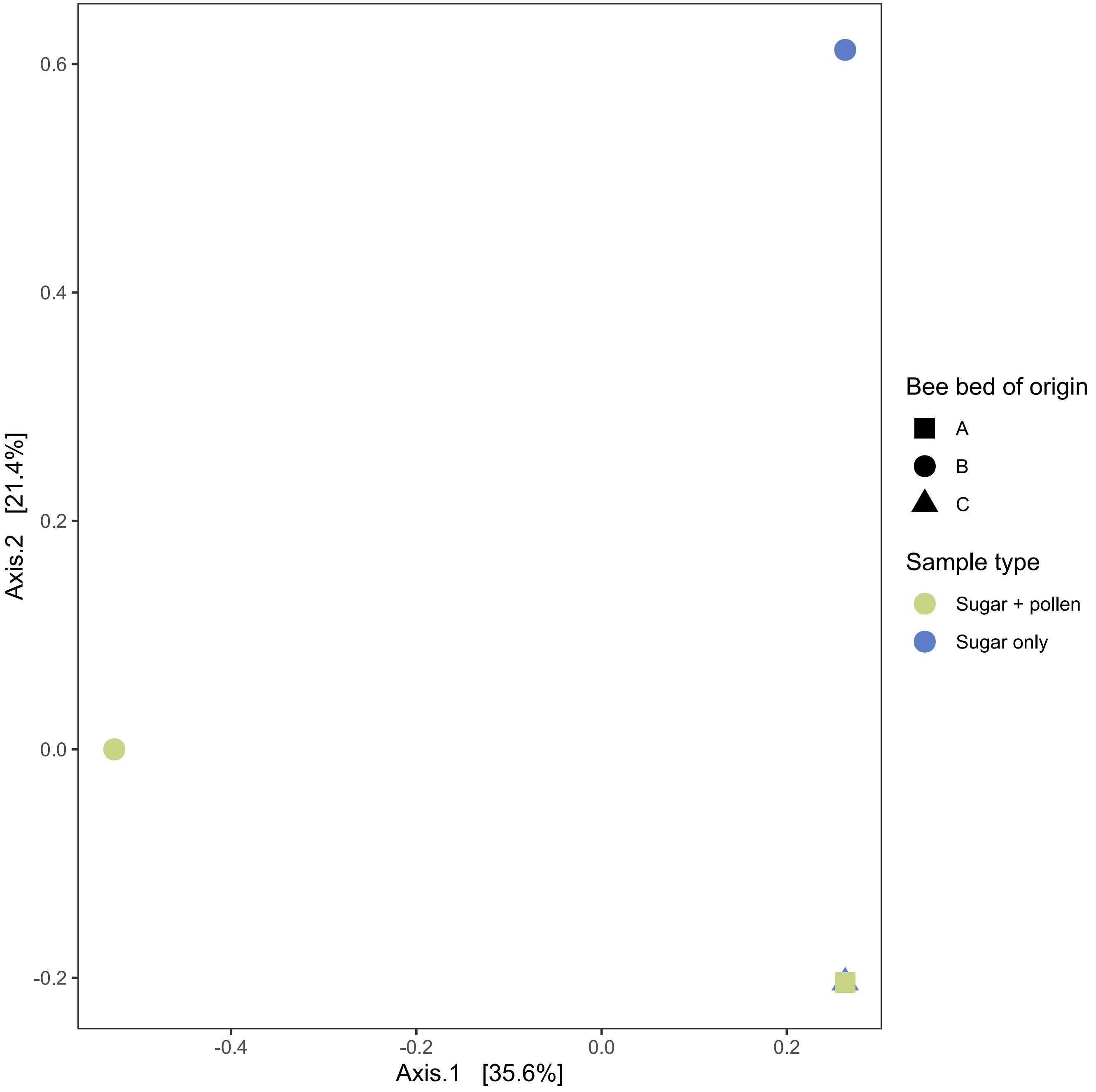


Fig. S1. Microbiome composition of lab-reared females. Principal Coordinates Analysis (PCoA) plot of Bray-Curtis dissimilarity. Each point represents that bacterial community of an individual sample. Colors indicate diet type and shapes indicate bee bed of origin.

**Fig. S2**

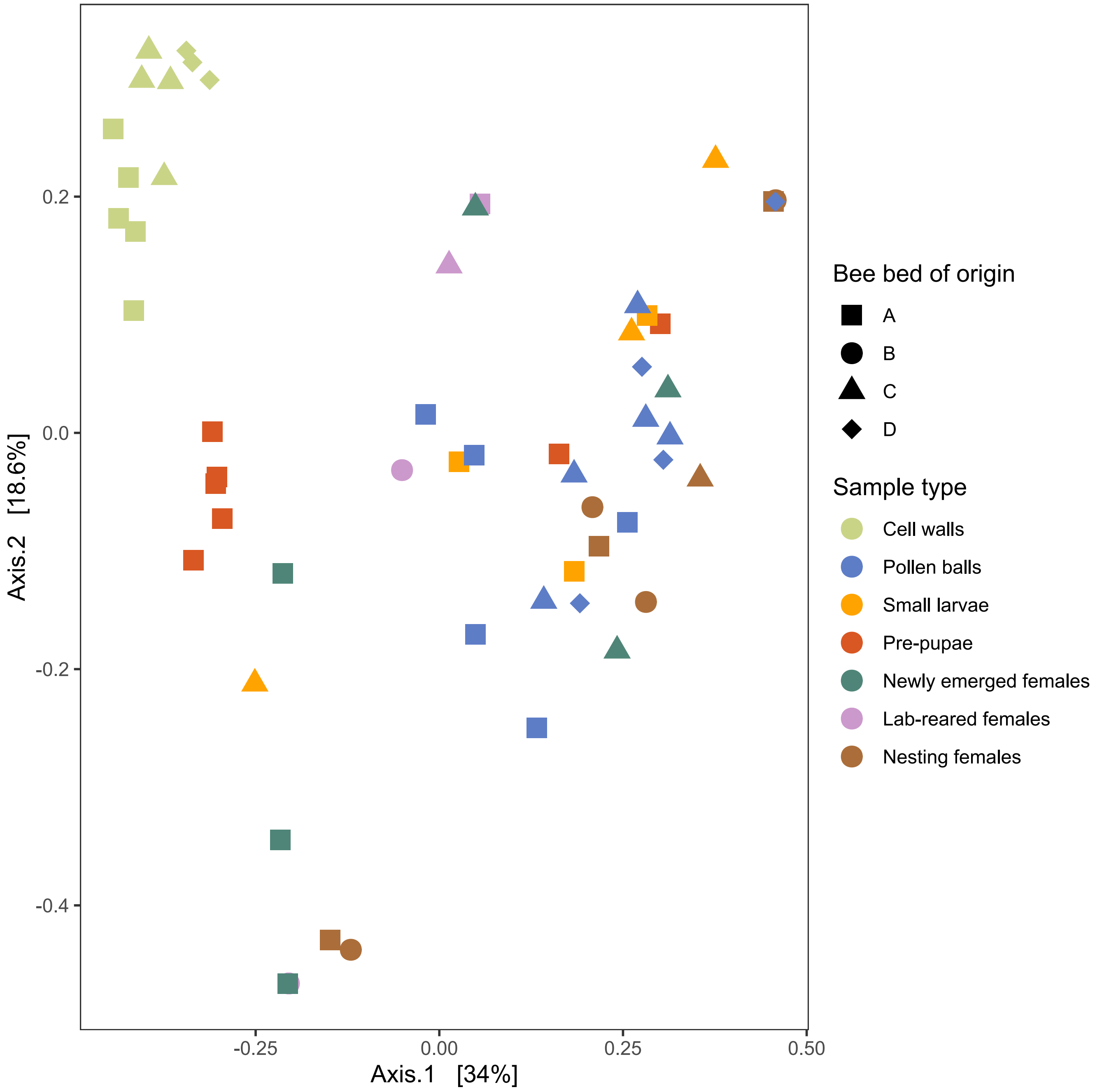


Fig. S2. Microbiome composition across the alkali bee life cycle. Principal Coordinates Analysis (PCoA) plot of weighted unifrac distance matrix. Each point represents the bacterial community of an individual sample. Colors indicate sample type and shapes indicate bee bed from which sample was collected.

**Fig. S3**

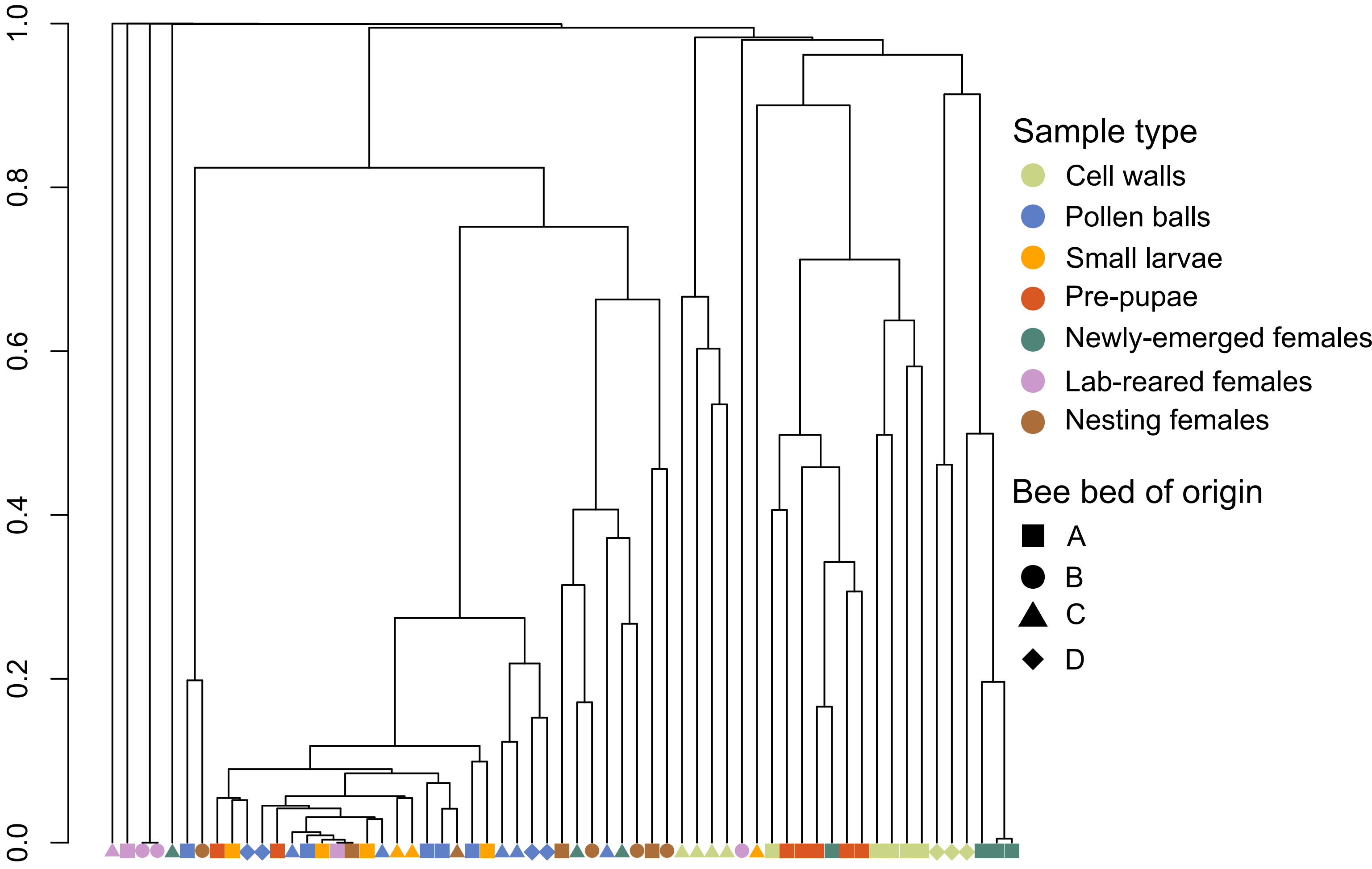


Fig. S3. Clustering of alkali bee samples based on average linkage of relative abundances within the microbiome. Each tip represents an individual sample. Colors indicate sample type and shapes indicate bee bed of origin.
